# Improved Metabolite Prediction Using Microbiome Data-Based Elastic Net Models

**DOI:** 10.1101/2021.07.01.450697

**Authors:** Jialiu Xie, Hunyong Cho, Bridget M. Lin, Malvika Pillai, Lara H. Heimisdottir, Dipankar Bandyopadhyay, Fei Zou, Jeffrey Roach, Kimon Divaris, Di Wu

**Affiliations:** Department of Biostatistics, Gillings School of Global Public Health, University of North Carolina at Chapel Hill, Chapel Hill, USA; Department of Genetics, School of Medicine, University of North Carolina at Chapel Hill, Chapel Hill, USA; Division of Oral & Craniofacial Health Sciences, School of Dentistry, University of North Carolina, Chapel Hill, NC, USA; Division of Pediatric and Public Health, Adams School of Dentistry, University of North Carolina, Chapel Hill, NC, USA; Department of Biostatistics, Virginia Commonwealth University, Richmond, VA, USA; Research Computing, University of North Carolina, Chapel Hill, NC, USA

**Keywords:** microbiome, metatranscriptome, metabolome, prediction, elastic net, random forest

## Abstract

Microbiome data are becoming increasingly available in large health cohorts yet metabolomics data are still scant. While many studies generate microbiome data, they lack matched metabolomics data or have considerable missing proportions of metabolites. Since metabolomics is key to understanding microbial and general biological activities, the possibility of imputing individual metabolites or inferring metabolomics pathways from microbial taxonomy or metagenomics is intriguing. Importantly, current metabolomics profiling methods such as the HMP Unified Metabolic Analysis Network (HUMAnN) have unknown accuracy and are limited in their ability to predict individual metabolites. To address this gap, we developed a novel metabolite prediction method, and we present its application and evaluation in an oral microbiome study. We developed ENVIM based on the Elastic Net Model (ENM) to predict metabolites using micorbiome data. ENVIM introduces an extra step to ENM to consider variable importance scores and thus achieve better prediction power. We investigate the metabolite prediction performance of ENVIM using metagenomic and metatranscriptomic data in a supragingival biofilm multi-omics dataset of 297 children ages 3-5 who were participants of a community-based study of early childhood oral health (ZOE 2.0) in North Carolina, United States. We further validate ENVIM in two additional publicly available multi-omics datasets generated from studies of gut health and vagina health. We select gene-family sets based on variable importance scores and modify the existing ENM strategy used in the MelonnPan prediction software to accommodate the unique features of microbiome and metabolome data. We evaluate metagenomic and metatranscriptomic predictors and compare the prediction performance of ENVIM to the standard ENM employed in MelonnPan. The newly-developed ENVIM method showed superior metabolite predictive accuracy than MelonnPan using metatranscriptomics data only, metagenomics data only, or both of these two. Both methods perform better prediction using gut or vagina microbiome data than using oral microbiome data for the samples’ corresponding metabolites. The top predictable compounds have been reported in all these three datasets from three different body sites. Enrichment of prediction some contributing species has been detected.

## INTRODUCTION

The importance of the human microbiome in health and disease is undeniable; site-specific microbial communities interact both with the environment and the host and influence numerous biological processes (1). Aside from the logical interest in understanding the microbiome’s composition, measuring and understanding its associated metabolic activities is arguably of utmost biological relevance. Recent studies have linked the metabolome with several important health conditions including inflammatory bowel disease (IBD)(2), obesity and type II diabetes (3), cholesterol levels (4), and early childhood dental caries (ECC)(5). Despite the rapidly increasing availability of microbiome data in large health cohorts, metabolomics data are still scant. This is an important limitation because the lack of, or considerable missingness of, metabolite information in microbiome studies can diminish their potential in inferring functions and important biological targets.

It follows that methods that help fill in the functional information gaps in microbiome studies are valuable and necessary. Because “matched” microbiome and metabolome datasets are extremely scant, most current methods rely on metabolic pathway inferences from taxonomic and metagenomic data, such as in the HMP Unified Metabolic Analysis Network (HUMAnN) (6). While the value of this approach is well-documented for the analysis of some microbial consortia (e.g., the human gut) (2, 7), HUMAnN cannot make predictions for individual metabolites. Moreover, its accuracy has not been benchmarked and its performance in other microbial communities with distinct ecology and function (e.g., the oral cavity) remains unknown. This is important because measured metabolomes at different body sites may include, besides the products of microbial metabolism, biochemical contributions from the host and the environment (e.g., dietary sugars in the study of dental biofilm(5)). Although an accurate determination of metabolite sources may not always be possible, predictions of these biofilm metabolites using microbiome information are highly desirable.

Along these lines, in 2016, Noecker and colleagues (8) added to the available analytical toolbox by leveraging 16S rRNA data. Their method enabled model-based integration of metabolite observations and species abundances using taxonomy and paired metabolomics data from ~70 vaginal samples. More recently, MelonnPan (9) was developed to obtain metabolomic profiling of microbial communities using amplicon or metagenomic sequences. This new method was motivated and applied in the context of paired microbiome and metabolome data in the context of an IBD cohort.

The motivation for the present new method development is to improve existing analytical approaches available for metabolite prediction and functions using microbiome data (10). To this end, we leverage existing microbiome and metabolome data from a study of early childhood oral health investigating ECC, a study of the human gut investigating IBD, and a study of vaginal health. The elastic net model (ENM, also used in MelonnPan), compared to LASSO or ridge regression, benefits from keeping both the singularities at the vertices, which is necessary to accommodate data sparsity, and the strict convex edges for grouping among correlated variables. Inspired by MelonnPan and MIMOSA, we propose an improved prediction method for individual metabolites using microbiome information in the same biological samples (as matched samples or paired samples), called “Elastic Net Variable Importance Model (ENVIM)”. It improves upon ENM algorithms by weighing microbial gene features using random forest variable importance (VI) to enhance the contribution of most prediction-informative genes. ENVIM outputs predicted metabolites from matched microbiome samples, as well as genes and their weights informing metabolite prediction.

In this paper, we present the development, application, and evaluation of the new method ENVIM. We compare it against MelonnPan in three datasets generated from oral, gut, and vaginal samples, so that we can also compare the metabolite prediction among different body sites. The preditors can be three different gene family data types, e.g., metagenome only, metatranscriptome only, and the combination of both metagenome and metatranscriptome data. The top predictable coumpounds have been reported in all these three datasets from three different body sites. Enrichment of some prediction contributing species has been detected.

## MATERIAL AND METHODS

### 1. Cohort and data description

In the following section, we describe the microbiome and metabolome data used for the new method development and application, alongside the three contributing studies.

#### ZOE 2.0 study data

ZOE 2.0 is a community-based molecular epidemiologic study of early childhood oral health in North Carolina (11, 12). The study collected clinical information on preschool-age children’s (ages 3-5) dental cavities (i.e., referred to early childhood caries or ECC) (13) and supragingival biofilm samples from a sample of over 6,000 children (14). A subset of participants’ biofilm samples underwent metagenomics, metatranscriptomics, and metabolomics analyses, under the umbrella Trans-Omics for Precision Dentistry and Early Childhood Caries or TOPDECC (accession: phs002232.v1.p1) (11). As such, metagenomics (i.e., shotgun whole genome sequencing or WGS), metatranscriptomics (i.e., RNA-seq), and global metabolomics data (i.e., ultra-performance liquid chromatography-tandem mass spectrometry) (5, 15, 16) from supragingival biofilm samples of ~300 children, paired with clinical information on ECC are available. After exclusions due to phenotype and metabolite missingness described in a previous publication (5), the joint microbiome-metabolome data include 289 participants. There are 503 known metabolites included in the ZOE 2.0 dataset. Metagenomics and metatranscriptomics data in reads per kilobase (RPK) were generated using HUMAnN 2.0. Here, we use species-level (205 species), gene-family (403K gene families), pathway (397 pathways), and metabolome (503 metabolites) data.

#### Lloyd-Price study data

The Lloyd-Price dataset (2) was obtained from the Inflammatory Bowel Disease multi-omics database (https://ibdmdb.org). It is derived from a longitudinal study that sought to generate profiles of different types of omics data among 132 participants for one year and up to 24 time points. The study’s several different types of omics data include WGS shotgun metagenomics, RNA-seq metatranscriptomics, and metabolomics. The corresponding metadata include demographic information such as occupation, education level, and age. These gut microbiome data are in counts per million (CPM) and were derived using functional profiles 3.0 in HUMAnN3.0. For this study, we merged data of individual gene families for 1638 samples for 130 subjects, and individual metatranscriptomics gene families for 817 samples for 109 subjects, respectively. The merged metagenomics gene families data include about 2,741K gene families and 1580 samples. Merged metatranscriptomics gene families data include about 1,079K gene families and 795 samples. The metabolomics data were generated using LC-MS and include 81,867 metabolites and 546 samples for 106 subjects. Most metabolites have not been annotated into known biochemicals and thus were excluded from prediction. After limiting the dataset to known metabolites and removing “redundant ions” in “HMDB” ID, there remained 526 metabolites to be predicted.

#### Mallick study data

The Mallick data (9) comprised the main real-life dataset used in the development of the MelonnPan method (9). They are derived from gut microbiome WGS shotgun sequencing from two cross-sectional IBD cohort studies, namely the Prospective Registry cohort for IBD Studies at the Massachusetts General Hospital (PRISM) and the Netherlands IBD cohort (NLIBD). Gene-family data in RPK units were derived using HUMAnN2.0 and normalized to reads per kilobase per million sample reads (RPKM). The raw metagenomics gene-family dataset includes 1 million gene families. The investigators (9) filtered out genes with low abundance and prevalence resulting in a processed dataset of 811 genes available in the R package *Melonnpan* (melonnpan.training.data and melonnpan.test.data) for 222 total subjects. The microbiome data have been pre-processed and normalized into relative abundance. The metabolite abundance data (8,848 metabolites and 220 subjects) have been made available by Franzosa et al (17). Those authors used 466 metabolites for analyses, a subset that was confirmed experimentally against laboratory standards prior to application in *Melonnpan*. In the present study, we use information from these 466 metabolites to compare the power of the new ENVIM method against *Melonnpan*. To accomplish this, we normalized the metabolite abundance data for all 8848 metabolites into relative abundance (compositional format). Among them, we used the same 466 metabolites with laboratory standards as selected in the paper of *Melonnpan* (9). Data missingness is not an issue in the Mallick metabolome data.

### 2. Metabolomics data pre-processing and normalization

An overview of the approach for metabolome data is presented in **Figure 1** and elaborated in detail below.

**Figure 1.**
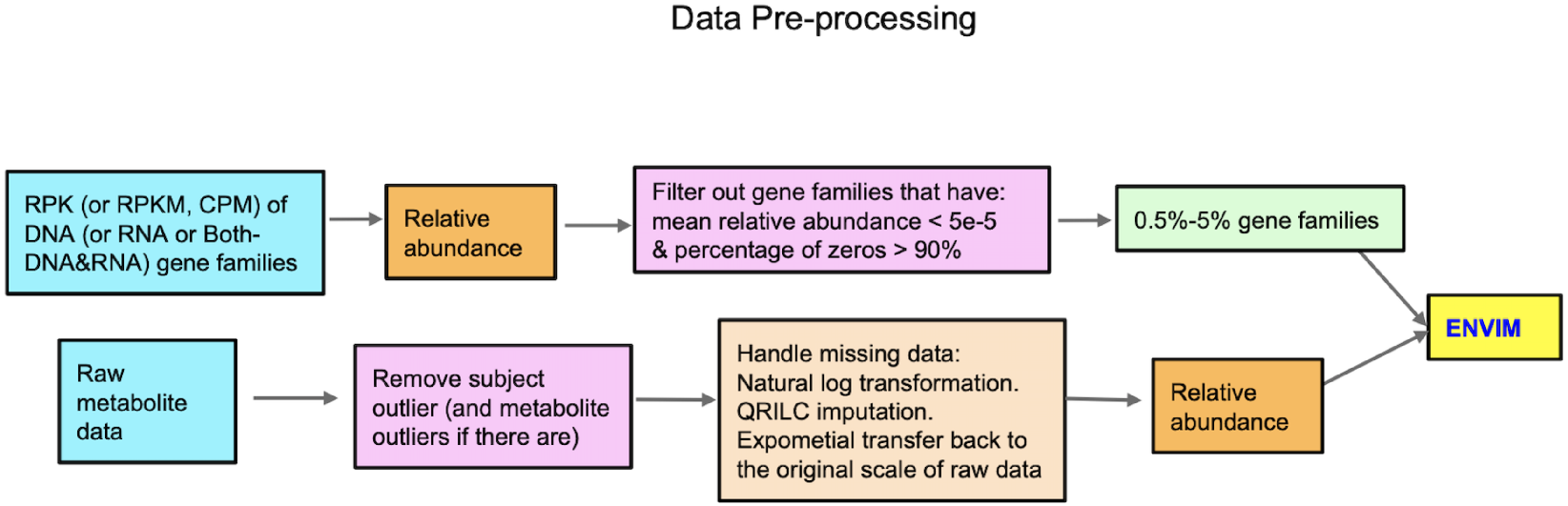
Flowchart of data preprocessing in microbiome and metabolome Mallick didn’t use QRILC, the other two used.

#### Metabolomics missing data imputation: ZOE 2.0 and Lloyd-Price studies

The proportions of missing metabolite data are small in the ZOE 2.0 and Lloyd-Price studies. In ZOE 2.0, 87% of metabolites have some missing data whereas 58% have missing values in Lloyd-Price. To address missingness in these two cohorts, we applied a rigorous feature-wise Quantile Regression Imputation of Left-Censored data (QRILC)(18) to impute missing metabolite values and avoid underestimated metabolite-level variance, as in a previous publication (5). All 503 metabolites in ZOE 2.0 have <90% missing data among the 289 included participants. We applied a similar preprocessing filter for the Lloyd-Price data (i.e., removing outlier subjects, **SuppFigure 1**), resulting in the exclusion of 15 outlier subjects with the largest numbers of missing metabolite values, as well as outlier metabolites with >90% missing values. Consequently, we carried forward to analyze 522 metabolites in 531 samples from the Lloyd-Price data.

The application of the QRILC imputation method departed from a natural log data transformation for the imputation step and an exponentiation to back transform the data to RPK (in ZOE 2.0) or CPM (in Lloyd-Price) scales. Because MelonnPan requires metabolite data to be inputted as compositional, we converted RPK and CPM imputed data to a compositional format before predictive modeling.

#### Metabolites Filtered by Metabolic Pathways (ZOE 2.0, Lloyd-Price and Mallick)

We used the MetaCyc database to retain only “reactive” metabolites. To achieve this, we considered metabolites’ membership in any MetaCyc metabolic pathway, reflecting reactions between bacteria and metabolites, and carried out the following steps:

1. In the MetaCyc database, we identify metabolites in each of the pathways predicted by metagenomics data in Functional Profile 2.0 generated by HUMAnN 2.0 (ZOE 2.0 and Mallick data) and Functional Profile 3.0 generated by HUMAnN 3.0 (Lloyd-Price data).
2. We used metabolites labels (KEGG id, HMDB, PUBCHEM, and metabolite name, provided in Metabolome data annotation, provided by manufacturer) in each of the three datasets, as the mapping IDs for each metabolite.
3. In MetaCyc, regardless of the metabolite label, only one unique MetaCyc “weblink” or universal mapping id is returned if the metabolite is in the database. This way, reactive metabolites identified in step 1 can be matched with metabolites identified in step 2. This way, we identify metabolites that are in the observed pathways. Finally, we filter out metabolites with low abundance (metabolites with mean relative abundance <10^−4^) and low prevalence (metabolites with >10% non-zero values). Consequently, there were 149 metabolites in pathways in ZOE 2.0, 125 in Lloyd-Price, and 251 in the Mallick data. Of note, no pathway information exists in the Mallick data. To compare the prediction of metabolites in pathways with the prediction of all metabolites, we considered both sets of metabolites in our analyses.

### 3. Microbiome data pre-processing and normalization

An overview of the approach for microbiome data is presented in **Figure 1** and elaborated in detail below. First, we matched gene family-level microbiome data with metabolome data by participant or sample unique identifier. Then, the scaled (RPK, RPKM, or CPM) gene family abundances were converted to compositional data, relative to the total per sample. Then, we filtered out gene family features with low relative abundance (mean relative abundance <5×10^−5^) and low prevalence (percentage of zeros in >90% of the samples) and thus kept 0.5% - 5% of gene family features. The same procedures were performed for both metatranscriptomics (briefly referred to as “RNA” thereafter) and metagenomics data (briefly referred to as “DNA” hereafter) thereafter, respectively. When both DNA and RNA data (briefly as “BOTH” hereafter) are considered predictors, a gene name may correspond to two “gene features”, one for each data type. The same data pre-processing and normalization procedures were followed for three cohorts, with sample sizes and feature numbers presented in **Table 1**. To prevent overfitting when evaluating ENM and ENVIM, we divided samples into training (75% of subjects) and testing datasets (25% of subjects).

**Table 1.** Sample size and number of selected gene family features. Testing genes: genes can be used in the testing set. Training genes: genes can be used in the training set. Genes in both: genes are in both training and testing sets.

|  |  | Training genes | Testing genes | Genes in both | Subjects | Metabolites | Metabolites (in pathways) |
| --- | --- | --- | --- | --- | --- | --- | --- |
| ZOE 2.0 | DNA (total 403k genes) | 1355 | 1276 | 1214 | 289 | 503 | 149 |
|  | RNA (total 403k genes) | 1805 | 1826 | 1667 | 287 | 503 | 149 |
|  | BOTH (total 806k genes) | 3158 | 3183 | 2948 | 287 | 503 | 149 |
| Lloyd-Price | DNA (total 2741k genes) | 726 | 712 | 633 | 359 | 522 | 125 |
|  | RNA (total 1079k genes) | 726 | 704 | 600 | 282 | 522 | 125 |
|  | BOTH (total 3820k genes) | 1424 | 1508 | 1211 | 269 | 522 | 125 |
| Mallick | DNA (total 1000k genes) | 811 | 811 | 811 | 220 | 466 | 251 (Filter Only) |

### 4. The existing ENM method for microbiome data-based metabolite prediction

As mentioned previously, the existing method available for predicting metabolite abundance using metagenomics data is MelonnPan(9) (Model-based Genomically Informed High-dimensional Predictor of Microbial Community Metabolic Profiles). In this study, in MelonnPan we used all filtered metagenomic gene family features in the 10-fold cross-validated elastic net model (ENM)(19) to predict metabolite abundance (**Equation 1**).

However, using all filtered metagenomic gene family features in the model may dilute the effect of some important gene family features contributing to the prediction of metabolite abundance. This limitation can be improved upon, and therefore, in this paper, we set out to improve the ENM and develop a new algorithm.

The MelonnPan software was downloaded from Github (https://github.com/biobakery/melonnpan) or in *Melonnpan* Package in R, the CSV output file “Predicted_Metabolites.txt” (Train) and “MelonnPan_Predicted_Metabolites.txt” (Test) are used as the prediction results of MelonnPan.

Elastic net model (ENM) assumes the model,

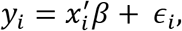

where *β* = (*β*_0_,*β*_1_,…, *β_p_*)′ and 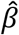, the ENM estimator of *β*, is found by minimizing the objective function of ENM,

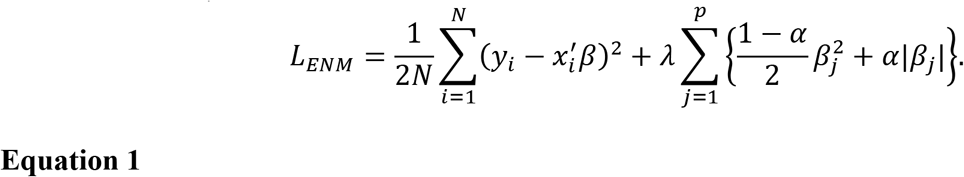

### 5. Evaluation Methods

We used Cohen’s criterion (20), to define Well-Predicted (WP) metabolites as those with Spearman correlation ≥0.3, and those with correlation <0.3 as poorly predicted. The predictive performance of the new method ENVIM is evaluated by comparing it against MelonnPan. Additionally, we compare Spearman correlations and mean square error (MSE) between the predicted and observed metabolites in both the training stage and the testing stage for all the three datasets and both methods.

## RESULTS

### 1. The improved ENM based on variable importance score (ENVIM)

The new algorithm is based on ENM, as the Elastic Net Variable Importance Model (ENVIM) (**Equation 2**). The strategy in ENVIM and the comparison between ENM and ENVIM are shown in **Figure 2**. Because ENM assumes that both independent and dependent variables follow a normal distribution, we rank-transform each gene family’s feature to a normal distribution by using the *rntransform* (21) function in the R package *GENABEL* for training data and testing data, respectively. The training metabolite abundance data are transformed to a normal distribution by Box-Cox transformation. After fitting the model in the training data, predicted metabolite abundances are transformed back to relative abundance with *λ* being determined by the training metabolite abundance data.

**Figure 2.**
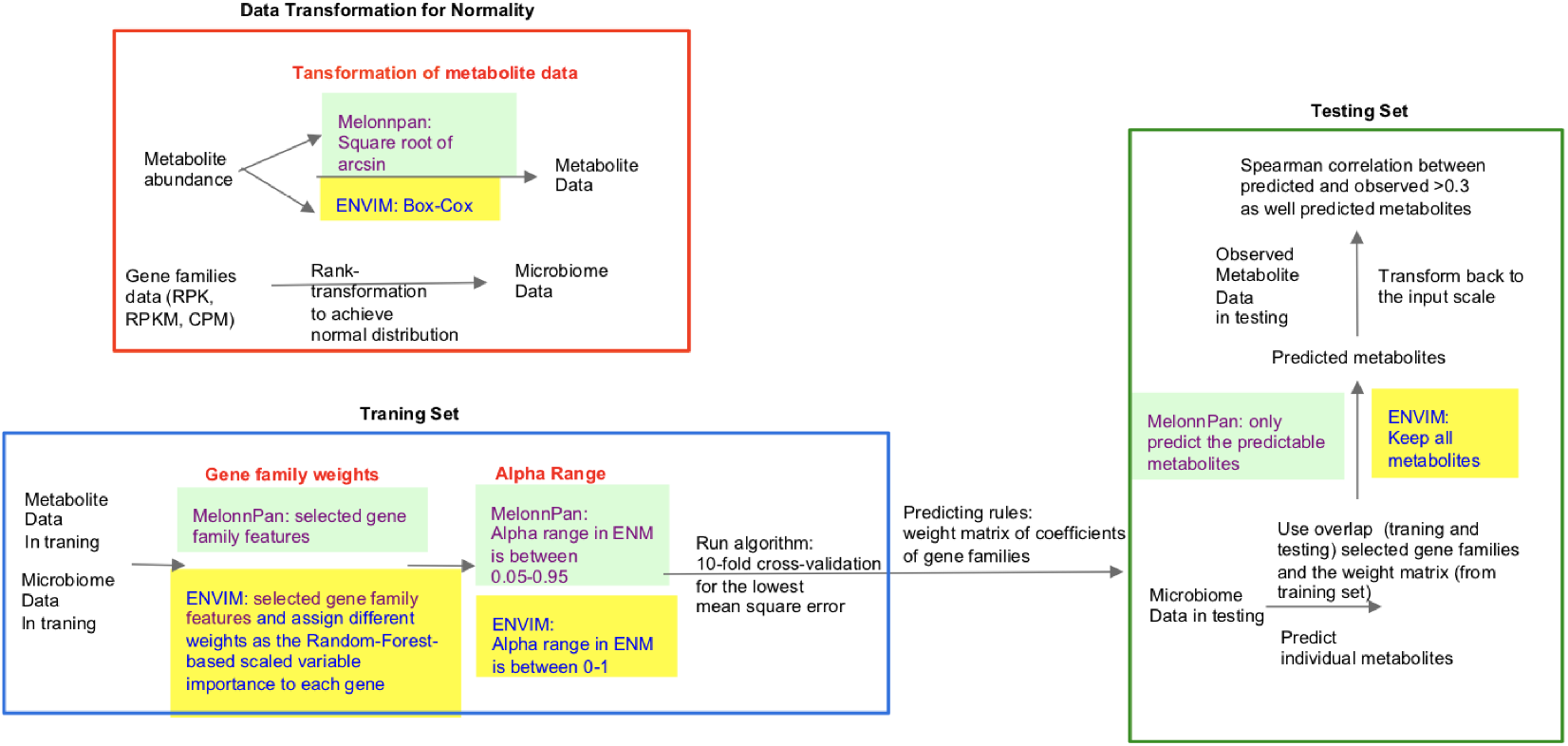
Flowchart of Melonnpan and ENVIM. The three differences between them include (red text) (1) Transformation of metabolite data (2) Gene family weights and (3) Penalty score. The predictable metabolites are defined as the metabolites that have a significant Spearman correlation with the adjusted q-value (testing whether the correlation is zero) below the default threshold in the training set.

Including all gene families into the model could make the cross-validated MSE larger, whereas including only a small part could make the error larger. Therefore, to identify a model with the minimum cross-validated error one needs to iterate different numbers of gene families. Because we prioritize gene families with high importance relative to metabolites, we use a nonlinear regression model to determine the importance of gene families for each metabolite. We train a cross-validated random forest model(22) by using the training data and use *varImp* function in the *caret* package(23) in R to find the scaled importance score (0-100) between each independent feature and the metabolite abundance. We introduce a unique step that uses the scaled variable importance score to classify gene families in different category intervals into the ENM, for example, gene families from 90 to 100. We use *glmnet* (24) package in R to run cross-validated ENM and choose penalty parameters for each model.

*In the training stage*, we divide the importance score from 0-100 into 10 category intervals (90-100, 80-100,…, 10-100, 0-100) and remove the intervals without gene families. We consider different sets of gene families with different importance scores into the ENM. Gene families are the independent variables, and metabolite abundances are the dependent variables. For each set of gene families, we conduct a 10-fold cross-validated ENM and build 10 models with different values of the tuning parameter *λ*, ranging from 0 to 1. For each model, we measure the MSE between the measured metabolite abundance and the predicted values to determine the best model (i.e., the model with the lowest MSE). To maintain reproducibility, we maintain the same random seed and permute the same fold index number in the ENM.

*In the testing stage*, for the prediction of each metabolite, we use a weight matrix of coefficients of gene families from the best model with the lowest MSE identified during the training stage. Because we previously transformed the compositional metabolite abundance data into compositional metabolite abundance per hundred and used Box-Cox transformation, we transform the predicted metabolite abundance data back to the original scale based on *λ* calculated in the training step. We evaluate the prediction of testing metabolite abundance by using the Spearman correlation between measured and predicted metabolite abundance data.

ENVIM assumes the following model:

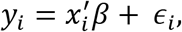

where *β* = (*β*_0_,*β*_1_,…,*β_p_*)′ and 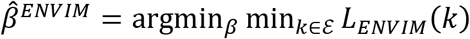, the ENVIM estimator of *β*, is found by minimizing over *k* and *β* the objective function,

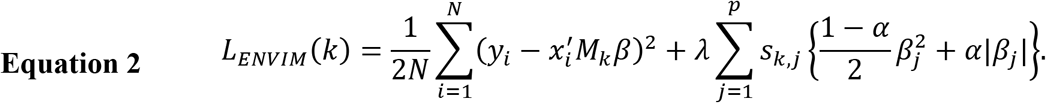

where we define *VI_j_* as the variable importance score for the *j*th variable given by a random forest, 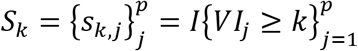 is the variable selection indicator vector giving 1 if the importance score for the jth variable is larger than the importance score *k*, 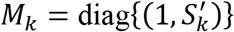 is the corresponding diagonal variable selection matrix that includes the intercept term, and *ε* is a set of the candidate *k* values. *ε* is defined adaptively so that it covers the range of the variable importance scores reasonably. In our analysis, we set *ε* ={0,10,20,…,90}.

**In the following sections, we present three key differences between MelonnPan and ENVIM for predicting individual metabolites (Figure 2).**

1. *Transformation of metabolite abundance data into a normal distribution*. To meet the assumption of ENM, MelonnPan transforms relative metabolite abundances with the arcsin square root operator, whereas we use Box-Cox transformation in ENVIM. To test normality, we compare the p-values of the Shapiro test statistics for both Box-Cox (**Equation 3**) and the arcsin square root transformations of metabolite abundances. In **Figure 3a**, the boxplots illustrate the distribution of -log10 of the p-values for all three data sets transformed by Box-Cox and arcsin square root and demonstrate that -log10 p-values for the Box-Cox transformation are greater than those applied with the arcsin square root transformation. In **Figure 3b**, in the scatter plot, -log10 p-values of almost all the metabolites applied with the Box-Cox transformation are more normally distributed than those applied with the arcsin square root transformation, implying that Box-Cox transformation yields better normal approximation than the arcsin square root transformation for most of the metabolites. Box-Cox transformation

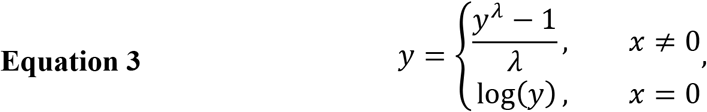

where *y* is the relative abundance.
2. *Different sets of gene families are carried forward to the prediction model*. MelonnPan uses all gene families in the training data in the ENM and ultimately predicts metabolites in the testing stage using the same features. However, regressing against all gene families may dilute the effect of important gene families. Thus, unlike MelonnPan, we use a variable importance criterion to select different sets of gene families and include them in the prediction models.
3. *α range in ENM* *Alpha* (*α*) is the weight between L_1_ and L_2_ penalty terms in the ENM, and in combination with *λ* values, the set of values that minimizes the 10-fold cross-validated MSE (Equation 1) is chosen. When *α* is 0, the model reduces to a ridge regression model which has the advantage of dealing with highly correlated independent variables; when *α* is 1, the model becomes a lasso regression model which has a variable selection capacity; when the *α* is between 0 to 1, the model includes the advantages of ridge regression and lasso regression. In MelonnPan, the range of *α* values does not include 0 and 1, which excludes either the Ridge or LASSO regression models, and it may not consider variables with high importance. The penalty term *alpha* in our ENVIM includes 0 and 1. By allowing a larger range of *α* we can include the case that is the Ridge regression model that does not exclude variables with high importance.

**Figure 3.**
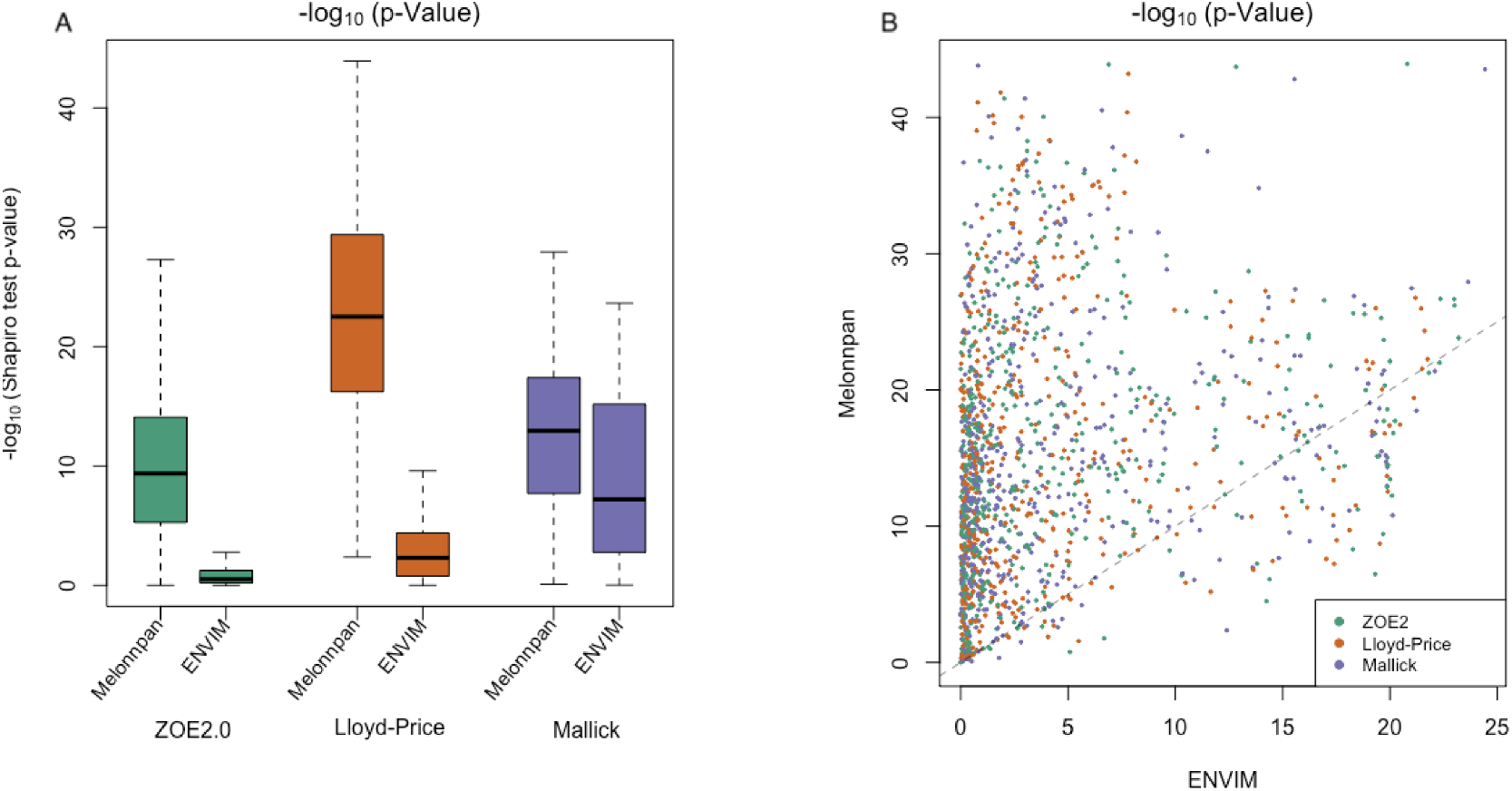
(a) Boxplot of -log10 of shapiro test p-values for relative metabolites abundances in all three data applied with box-cox transformation (we used) and arcsin square root transformation (Melonnpan used). (b) Scatter plot for comparing -log10 of p-values made by shapiro test (normality) between box-cox transformation (x-axis) and arcsin sqrt (y-axis) transformation. Almost all of the points are above the y = x line, which means that the -log10 of p-value after box-cox transformation is smaller than after arcsin sqrt transformation, and normality after box-cox transformation is better.

The ENVIM software written in R statistical language is available in Github (https://github.com/jialiux22/ENVIM). The “ENVIM_predict” function is for metabolite prediction only, and the ENVIM function is for metabolite prediction and evaluation given metabolomics data in the testing set is also available. Both will output weight matrix. The weight matrix in testing has the same values as in training if they have the same number of genes. Usually, testing has a smaller number of genes to be used for prediction, so the weight matrix in testing can be a subset of the weight matrix in the training set due to the smaller number of genes in the testing set.

### 2. Method Comparison for Prediction of Individual Metabolites in Three Datasets

#### Correlation-based method comparison

We used microbial gene family data to predict the individual metabolites in the matched samples (that are from the same biological sample in that one proportion is for microbiome and the other is for metabolome). We compared the prediction results between ENVIM and MelonnPan, in terms of Spearman correlation and mean square error (MSE) between predicted and observed values of each of the filtered metabolites, in three datasets (ZOE 2.0, Mallick data, and Lloyd-Price data) at each of the three data types of microbial gene families as the DNAseq, RNAseq, and BOTH (of RNA and DNA). MSE in the testing set is for comparison between methods (**Supplemental Figure 2**).

We have summarized the prediction results (**Table 2, Figure 4**) for all metabolites in terms of Spearman’s correlation according to three aspects: method comparison, modality comparison, and microbial community (i.e., body site) comparison. Overall, ENVIM produces higher percentages of well-predicted metabolites than MelonnPan in all three data cohorts, in both testing and training sets, and for DNA, RNA, and BOTH when available (**Table 2**).

**Table 2.** Prediction results (first four columns of numbers) in terms of Spearman correlation for all metabolites to be predicated. Based on the “well-prediction” criterion, defined as Spearman correlation > 0.3 between the observed and the predicted metabolites, the numbers of well-predicted metabolites with different prediction methods, datasets, and modality levels (DNA, RNA, and BOTH), are presented for comparing MelonnPan and ENVIM. NM is the number of metabolites to be predicted. Percentages in parentheses (%) represent the number of well-predicted metabolites divided by the total number of metabolites (NM) to be predicted in each study. The Mallick cohort has only metagenomics data available. The last column of numbers is for the numbers of “predicable metabolites”, that are defined by MelonnPan, also seen in the **Figure 2** legend.

| ZOE 2.0<br>(NM=503) | Training<br>(ENVIM) | Training<br>(MelonnPan) | Testing<br>(ENVIM) | Testing<br>(MelonnPan) | Predictable<br>metabolites<br>(Defined by<br>MelonnPan) |
| --- | --- | --- | --- | --- | --- |
| DNA only | 356 (71%) | 63 (13%) | **124**<br>**(25%)** | 47 (9%) | 70 |
| RNA only | 409 (81%) | 157 (31%) | 106 (21%) | 68 (14%) | 163 |
| BOTH DNA and<br>RNA | 423 (84%) | 146 (29%) | 110 (22%) | 73 (15%) | 154 |
| Mallick Cohort<br>(NM=466) |  |  |  |  |  |
| DNA only | 408 (88%) | 239 (51%) | 225 (48%) | 178 (38%) | 249 |
| Lloyd-Price<br>Cohort<br>(NM=522) |  |  |  |  |  |
| DNA only | 501<br>(96%) | 271 (52%) | 322<br>(62%) | 193 (37%) | 305 |
| RNA only | 521<br>(100%) | 298 (57%) | **393**<br>**(75%)** | 236 (45%) | 318 |
| BOTH DNA and<br>RNA | 518<br>(99%) | 306 (59%) | 381<br>(73%) | 232 (44%) | 323 |

**Figure 4.**
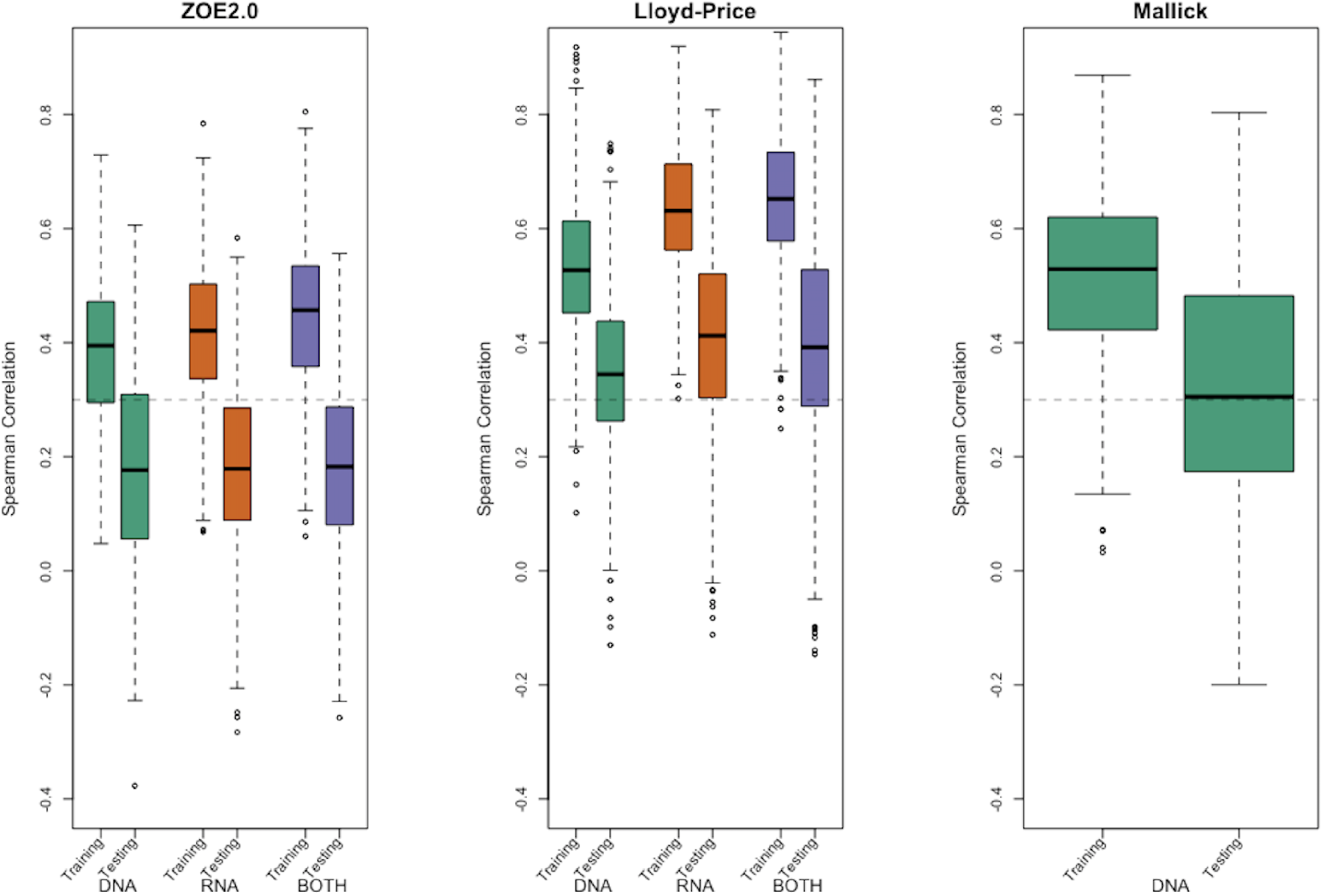
Evaluation using Spearman correlation in training stage and testing stage between predicted values and the observed values by using DNAseq data only, RNAseq data only, and both for ZOE2.0 data, Lloyd-Price Data, and Mallick data.

In general, RNA gene family data produce higher percentages of well-predicted metabolites than DNA data. In Lloyd-Price data, RNA-only data typically give higher percentages of well-predicted metabolites. In ZOE 2.0 and Lloyd-Price data, both DNA and RNA predictors produce similar percentages but are not always superior to the DNA-only or RNA-only data-based predictors. However, results emanating from both DNA and RNA predictors are never the worst. Not surprisingly, the well-predicted percentage of metabolites in testing sets is lower than in the training set (**Table 2**). The boxplots of Spearman correlations between the predicted and observed metabolites for all metabolites (**Figure 4**) show the overall distribution of Spearman correlation and suggest that the correlation between the ENVIM-predicted and the observed metabolites is higher in RNA than in DNA, but slightly lower than in both DNA and RNA. In testing, MelonnPan only predicts the predictable metabolites (defined as Well-Predicted metabolites in Training set, last columns in **Table 2**); while it is not appropriate to compare the correlation distribution for all metabolites as in **Figure 4** for MelonnPan. When comparing the distribution of correlation (**Figure 4**) between the ENVIM-predicted and the observed metabolites in ZOE2.0 and Lloyd-Price, the combination of both DNA and RNA appears to have higher correlations than the DNA only or RNA only in the training set. In testing, RNA data produce the highest median correlation. All three gene family data result in similar correlations in ZOE 2.0. It must also be noted that the highest proportion of well-predicted metabolites is found in the gut microbiome (Lloyd-Price) study, then in the vagina microbiome (Mallick study), and the lowest was in the supragingival dental biofilm (ZOE 2.0 study) (**Table 2**). Because in both the Lloyd-Price and Mallick datasets prediction correlations are higher than in ZOE 2.0 (**Figure 4**), it is reasonable to suggest better metabolite prediction in these sites and microbial communities than in the oral cavity.

Besides comparing MelonnPan and ENVIM in terms of percentages of well-predicted metabolites, one can directly compare the correlations of each predictable metabolite that is predicted by both methods (**Figures 5 and 6**). In the training set (**Figure 5**), all DNA, RNA, BOTH DNA, and RNA, and in all three datasets, we find that the majority of these metabolites have higher correlations in ENVIM compared to MelonnPan. The same holds in the testing set (**Figure 6**): most points are along the diagonal line but slightly above it, suggesting that metabolites predicted by ENVIM have higher correlations with the observed ones compared to those predicted by MelonnPan. We also find that there are more metabolites in the “ENVIM>0.3” category (blue) than in the “MelonnPan>0.3” category (red). This is a reflection of more well-predicted metabolites found after ENVIM than after MelonnPan prediction.

**Figure 5.**
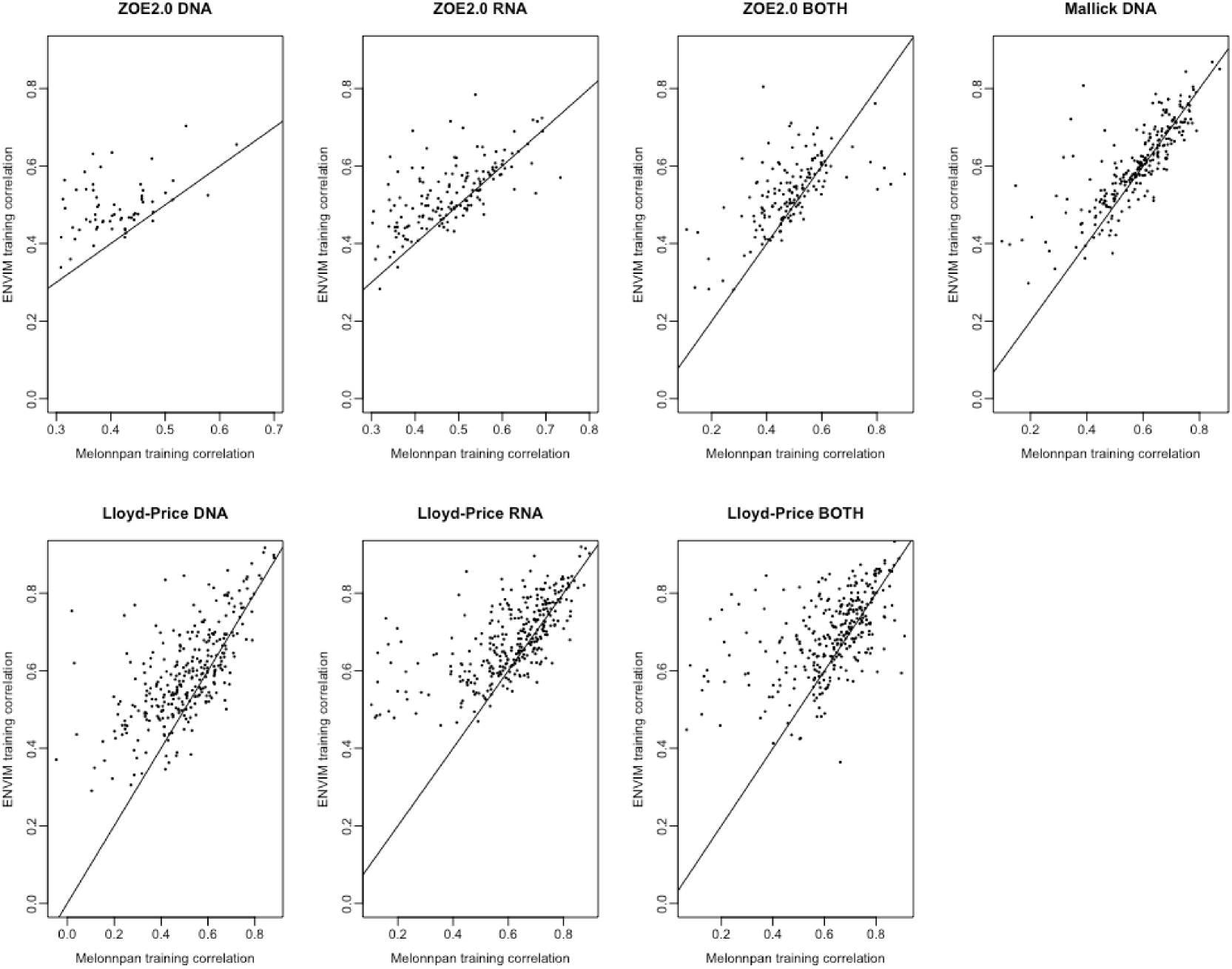
For DNA, RNA, and both in each study and the training set, this shows the scatter plot of Spearman correlation in ENVIM (y-axis) and Melonnpan (x-axis). Spearman correlation is based on observed metabolite abundance and predicted values. If our calculated correlation is NA, the metabolites will be included in this figure.

**Figure 6.**
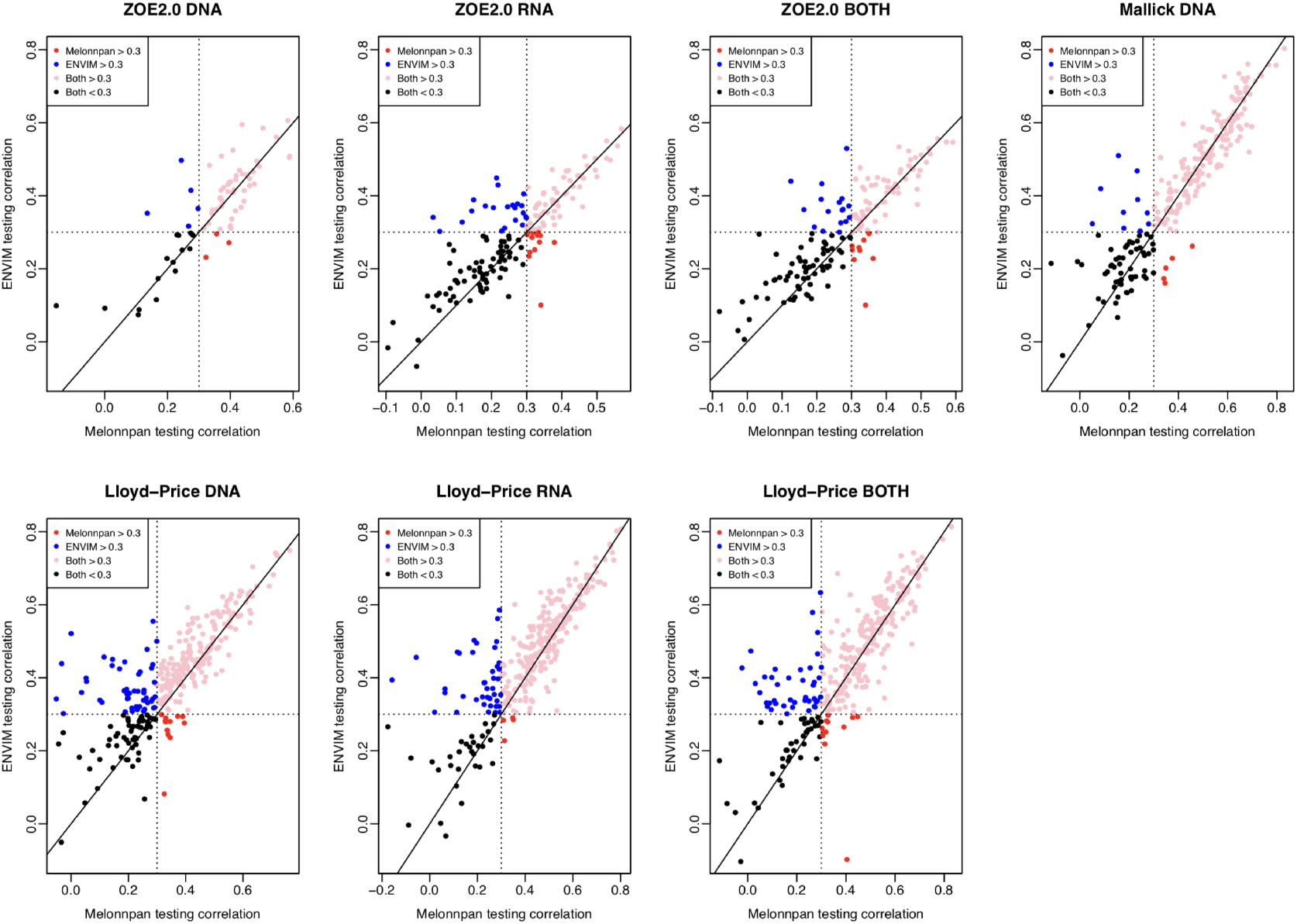
For DNA, RNA, and both in each study and the testing set, this shows the scatter plot of Spearman correlation in ENVIM (y-axis) and Melonnpan (x-axis). Spearman correlation is based on observed metabolite abundance and predicted values.

### 3. Methods comparison for prediction of individual metabolites in three datasets and the context of observed metabolic pathways

Metabolites may be associated with the microbiome in the context of metabolic pathways that involve interactions between host, microbiome, and environment. We further test the methods’ predictive power for metabolites that are found in microbiome data-based metabolic pathways generated by metagenomics and metatranscriptomics analysis in HUMANN2. Gene families are filtered by metabolic pathways as previously described in the methods section. All conclusions regarding the prediction of metabolites still hold in this scenario. Additionally, when comparing the percentages of well-predicted metabolites among all metabolites (first four columns of **Table 2**) and the metabolites found in pathways (**Table 3**), we find higher predicted percentages for the latter.

**Table 3.**
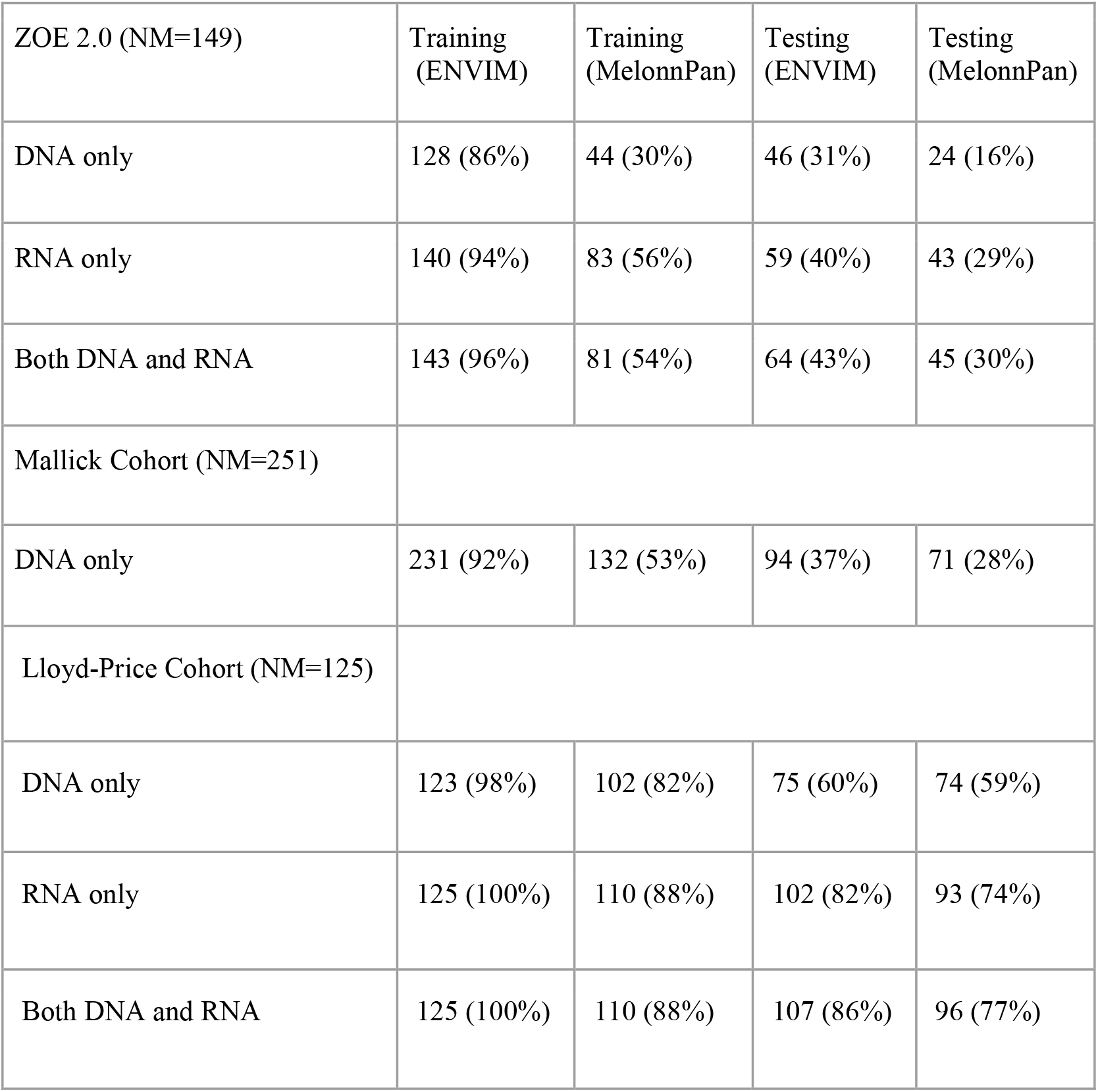
Prediction results via Spearman correlation for metabolites that are found in metabolic pathways. Based on the criterion of Spearman correlation >0.3 between observed and predicted metabolites, we present the numbers of well-predicted metabolites with different prediction methods, datasets, and modality levels (DNA, RNA, and both), and comparing between MelonnPan and ENVIM. NM is the number of metabolites to be predicted. Percentages in parentheses (%) represent the numbers of well-predicted metabolites divided by the total number of metabolites (NM) to be predicted in each study. The Mallick cohort has only metagenomics (DNA) data available and no pathway RNA data. The results from the Mallick cohort here are only based on filters (filtering out metabolites with mean relative abundance <10^−4^) and low prevalence (metabolites with >10% non-zero). In ZOE 2.0 and Lloyd-Price, metabolite data presented in this table have been selected according to membership in pathways and also satisfy the above-mentioned filtering criteria.

| ZOE 2.0 (NM=149) | Training<br>(ENVIM) | Training<br>(MelonnPan) | Testing<br>(ENVIM) | Testing<br>(MelonnPan) |
| --- | --- | --- | --- | --- |
| DNA only | 128 (86%) | 44 (30%) | 46 (31%) | 24 (16%) |
| RNA only | 140 (94%) | 83 (56%) | 59 (40%) | 43 (29%) |
| Both DNA and RNA | 143 (96%) | 81 (54%) | 64 (43%) | 45 (30%) |
| Mallick Cohort (NM=251) |  |  |  |  |
| DNA only | 231 (92%) | 132 (53%) | 94 (37%) | 71 (28%) |
| Lloyd-Price Cohort (NM=125) |  |  |  |  |
| DNA only | 123 (98%) | 102 (82%) | 75 (60%) | 74 (59%) |
| RNA only | 125 (100%) | 110 (88%) | 102 (82%) | 93 (74%) |
| Both DNA and RNA | 125 (100%) | 110 (88%) | 107 (86%) | 96 (77%) |

### 4. Methods comparison based on MSE

We use boxplots to compare the mean square errors (MSE) between measured and predicted metabolite abundance between ENVIM and MelonnPan both for training and testing models, with application to training data and testing data for all three studies. We only compare well-predicted metabolites identified by MelonnPan in training because MelonnPan only generates results for these metabolites. The boxplot demonstrates that the distribution of MSE in the MelonnPan model is approximately the same as the distribution of MSE in ENVIM (**Supplemental Figures 2**). There is no significant MSE difference between ENVIM and MelonnPan suggesting that both models predict these metabolites well, but the advantage of ENVIM is that we can predict substantially more well-predicted metabolites than MelonnPan—a consequence of MelonnPan’s inability to build a well-performing model in the training step.

### 5. Prediction Results of Individual Metabolites and Gene Weights in ENVIM

The top 50 predicted metabolite compounds from ENVIM across three datasets are shown in **Figure 7**. For Lloyd-Price and ZOE 2.0, we choose the gene family data that has the best ENVIM prediction power to show their top predicted metabolites, which are the DNA gene family data (124 metabolites as 25% among NM, **Table 2**) in ZOE 2.0, and the RNA gene family data (393 metabolites as 75% among NM, **Table 2**) in Lloyd-Price. The Mallick study only has DNA data available for metabolite prediction. Lloyd-Price data and Mallick data have measured metabolites in >1 metabolome LC-MS flatforms (see Data Description Section) so that one metabolite may appear >1 time in the top list, for example, the metabolite so-called Urobilin appeared in the top 50 for >1 time.

**Figure 7.**
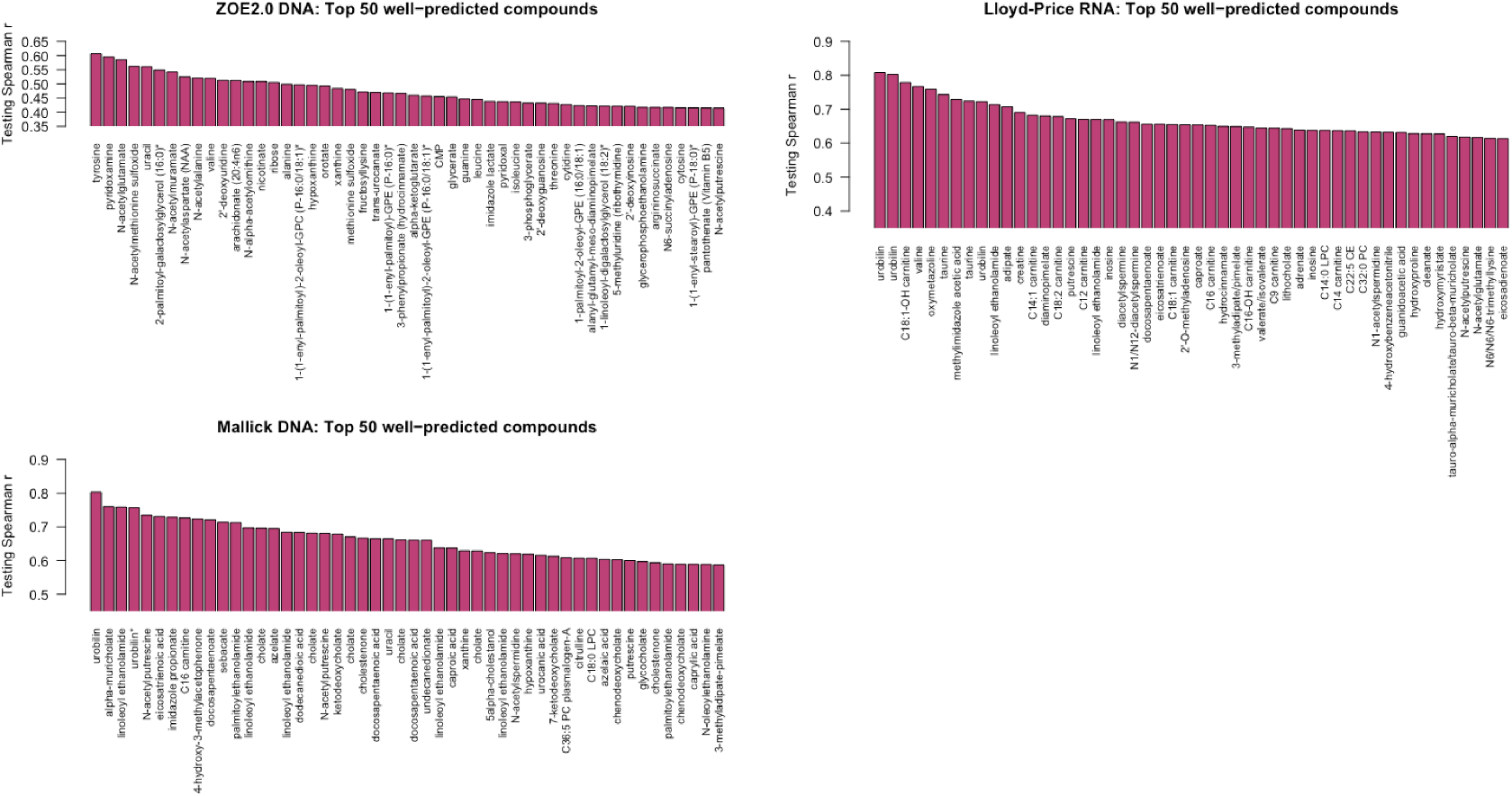
The best predicted 50 metabolite compounds (x-axis) in the three studies by ENVIM in the testing set. For Lloyd-Price and ZOE2.0, we choose the gene family data types that have the best ENVIM prediction power to show their top predicted metabolites, based on **Table 2**.

The summarized prediction results can be seen in **Supplemental Table 1**. To interpret the results, we take the carbohydrate pathway as an example that may provide the bacteria nutrition, so a few compounds have been well-predicted by the RNA gene data. We are aware the prediction in this paper is not about longitudinal causal relation but for mathematical prediction. Here we show four examples (**Figure 8A, B, C, D** for Trehalose, Maltose, Ribose Stachyose) that also have high Spearman correlation in the log10 scale of compositional data.

**Figure 8.**
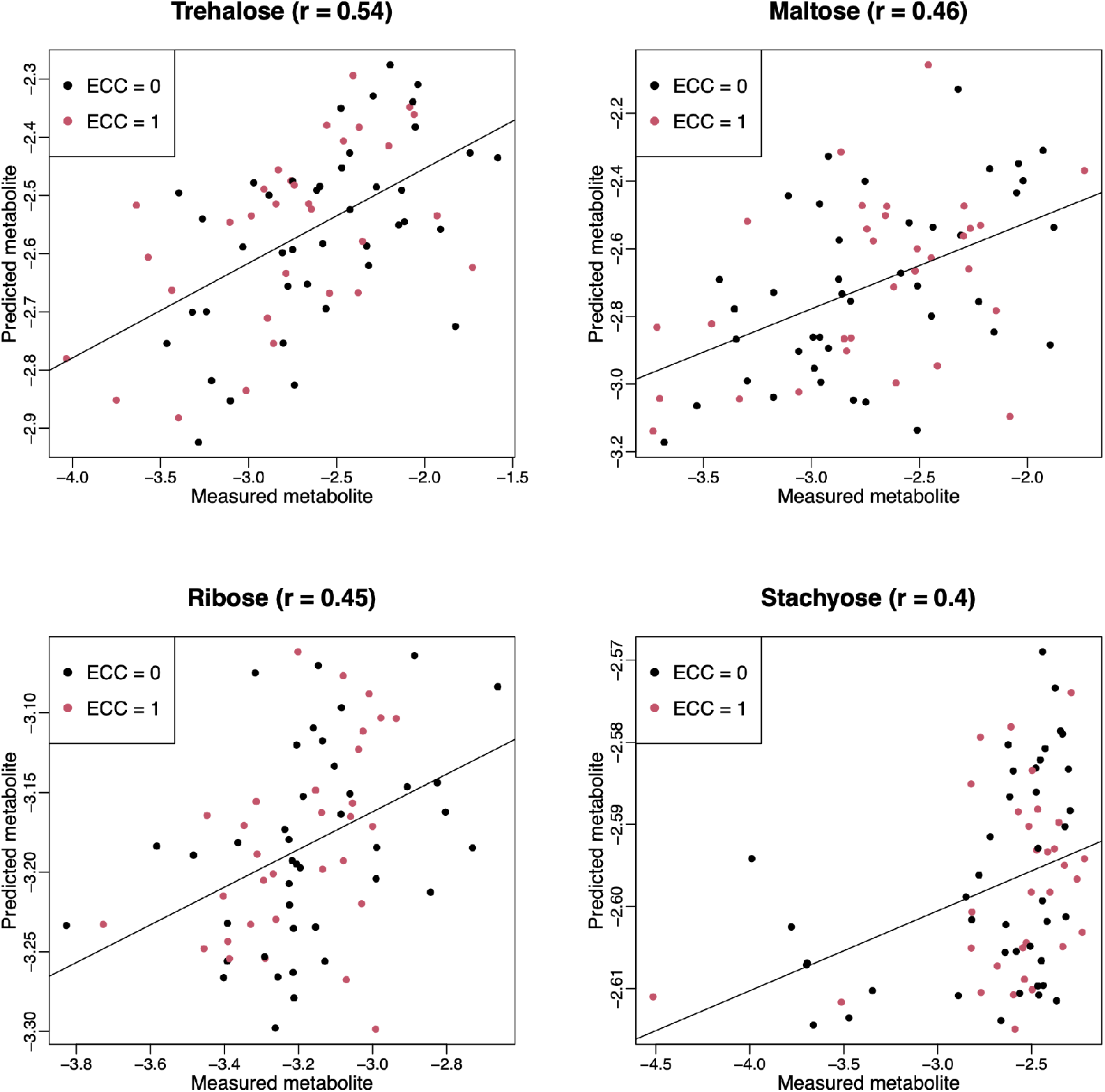
Scatter plots of examples of well-predicted metabolites in ZOE 2.0 by ENVIM. The X-axis is observed metabolites; the y-axis is for predicted metabolites. Both are in log10 scale of the compositional data for normality. ECC is for Early Childhood Caries, ECC =0 (about 50% of total samples in ZOE 2.0) is for the healthy group, and ECC=1 (about 50% of total samples in ZOE 2.0) is for the ECC case group. r is for Spearman correlation.

#### *Gene list (Weight matrix) comparison across three datasets, in ENVIM* (Supplemental Table 2)

We extract gene names that are non-zero in the weight matrix for each metabolite, dataset, and data type. We aim to compare gene names among three datasets and find the probability of predicting metabolites by using a different dataset. We find that there are not many overlapped genes (n < 10) between ZOE 2.0 data and Lloyd-Price data (Data not shown).

#### Gene set enrichment analysis (GSEA) within Species in ZOE 2.0

We perform gene set enrichment analysis to find the over-represented species of the gene families when we build the prediction model on metabolite abundance. We extract the weight matrix, merge the important gene families with non-zero values among all well-predicted metabolites. We get the summation of the rank of each gene family in the weight matrix based on the absolute value of the coefficient for each gene family. We use gene families data at the species level to find the species corresponding to those important gene families. For each species of bacteria, we compare the general difference in the cumulative distributions of gene families’ rank scores between each species and background species and find Kolmogorov–Smirnov (KS) p-values. We use the Benjamini–Hochberg false discovery rate (FDR) approach to correct the p-values and get q-values. There are 36 species in ZOE 2.0 DNA data and 73 species of bacteria in ZOE 2.0 RNA data that show significantly (q<0.05) over-represented species during the gene set enrichment analysis (**Figure 9**).

**Figure 9.**
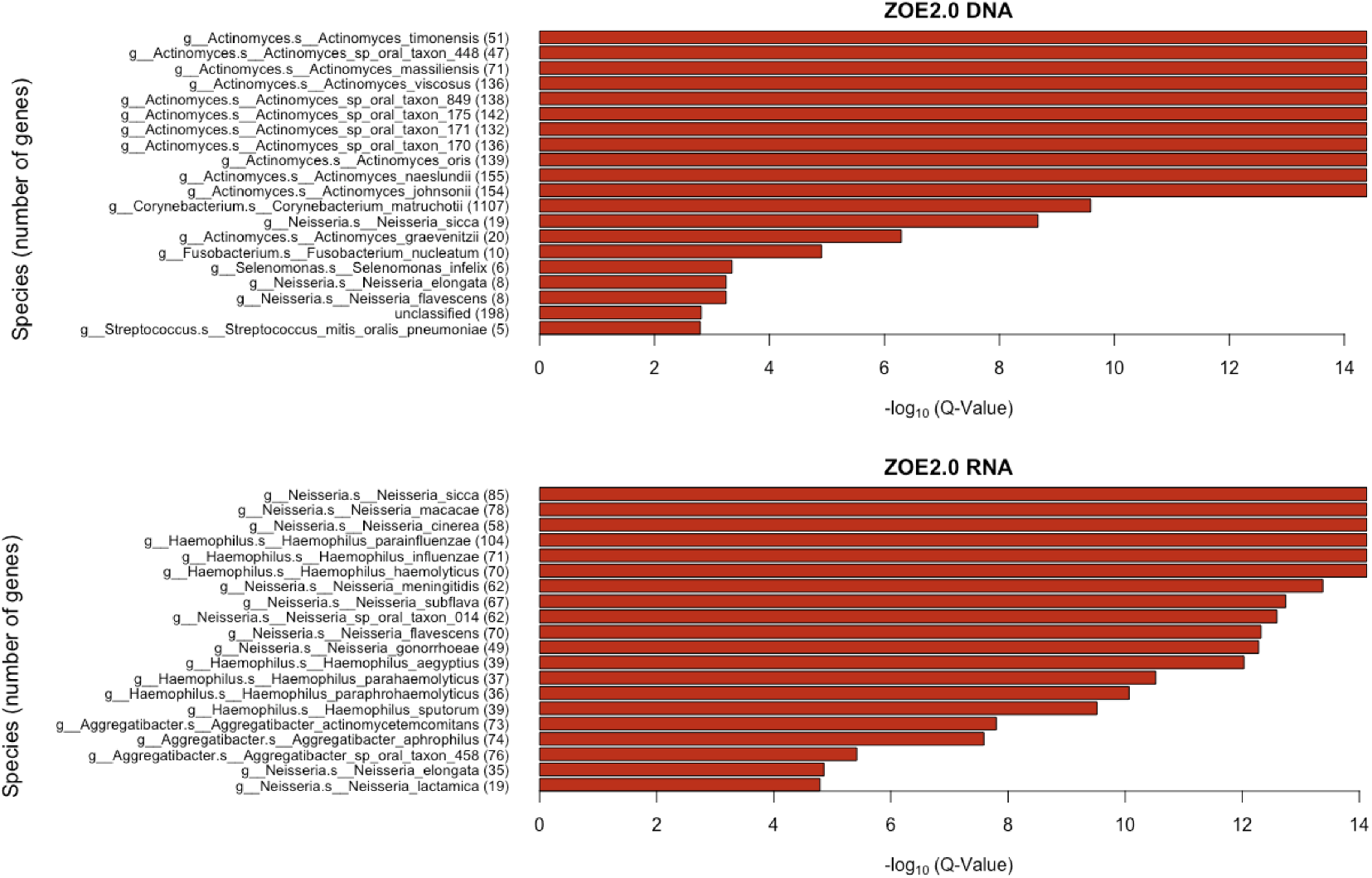
Taxonomic enrichment of metabolite predictive species for the most contributing species to metabolite prediction, based on ZOE2.0 DNA or RNA by ENVIM. The top 20 significant over-represented bacteria with the smallest Q values (Q < 0.05) for ZOE 2.0 data. The Q-value is based on the Kolmogorov-Smirnov (KS) test p values after FDR correction. (a) DNA data (b) RNA data.

Here, we used a different procedure for the gene set enrichment tests compared to what MelonnPan (9) used. They pooled genes in genera instead of species, due to the small number of genes in each species in their prediction procedure. We keep many more genes than MelonnPan so that we can address the ranks of genes instead of the binary prediction power of genes (i.e., whether a gene is used for prediction or not). Our GSEA strategy also can help avoid the bias to pick up the species that have larger numbers of genes.

### 6. Computational speed (compare to others)

Our developed method of improving ENM could run on the software on R and accurately predict metabolites. The mean prediction time for each metabolite for DNA gene families data is 5.2 minutes for ZOE2 data (6.1 minutes for Lloyd-Price Data, 2 minutes for Mallick data); mean prediction time for RNA gene families data is 4.2 minutes for ZOE2 data (3.7 minutes for Lloyd-Price Data); mean prediction time for both DNA and RNA gene families data is 4.5 minutes for ZOE2 data (3.6 minutes for Lloyd-Price Data) with MacOS Big Sur Version 11.4.

## DISCUSSION

We propose a new computational method for metabolite prediction using microbiome data-based improved Elastic Net Models. We chose different gene-family sets based on SVM-based variable importance scores and modified the existing ENM to accommodate the unique features of microbiome and metabolome data. The newly developed method ENVIM predicts metabolites using metagenomics, metatranscriptomics, or both data types. We apply the algorithm in three datasets, i.e., ZOE 2.0, Mallick, and Lloyd-Price studies. These three studies are mostly all we can find that have both microbiome and metabolome data in the same matched samples, with reasonably large sample sizes. Our work is the first time that researchers can use microbiome data to predict metabolites in more than one study, and different body sites. In addition, ZOE 2.0 and Lloyd-Price studies have both metagenomics and metatranscriptomics, so that we can for the first time, compare the prediction performance using the different gene family modalities (or called data types).

We evaluated metagenomic and metatranscriptomic predictors and compared the prediction performance between the previously developed MelonnPan and ENVIM, among DNA, RNA, and Both DNA and RNA gene families data using (1) the proportion of “well-predicted” metabolites defined as those with Spearman correlation between measured and predicted metabolite values > 0.3, (2) distribution of Spearman correlation and (3) MSE. The correlation suggests Both (using DNA and RNA jointly) provides robust prediction results that are never the worst among the three data types. Whether DNA or RNA have better prediction performance depends on the studies. The percentage of well-predicted metabolites is higher for metabolites that are in a metabolic pathway that is observed in microbiome data, and this supports the interaction between microbiome and metabolites may highly be related in known metabolic pathways. Across all datasets and data types, with or without the pathway filter, we find ENVIM always outperforms MelonnPan. We also find the prediction performance is better in Lloyd-Price and Mallick than in ZOE 2.0, which may suggest the association between microbiome and metabolites are stronger in the gut than in the oral cavity since oral metabolites may be more affected by environmental factors like food intake. More microbial omics studies are needed to compare the prediction power across different body sites and to understand how microbiome interact with metabolites differently at different body sites.

We are aware the data-preprocessing step has larger effects on the prediction performance. The distribution assumption, normalization, transformation, outlier filtering, and how to handle missing data are important to be considered before performing prediction. We have touched base on that, but potential further exploration may be needed.

The numbers of the measured metabolites and the numbers of the to-be predicted metabolites in each of the three studies are very different due to the difference of the technology platforms, and the available data. As what we proposed is not for causality but for mathematic prediction, we show examples of four metabolites (**Figure 8**) that may provide nutrition to species.

As a limitation, same as MelonnPan, the experimental design hasn’t been considered in this framework for ENVIM, including time course or disease statuses. As the purpose of this study is prediction, it’s reasonable to think that prediction is not necessary to be conditional on the experimental design. Instead, different disease statuses may have different microbiome profiles and have corresponding different metabolomes. Therefore, this is a limitation but not a drawback of prediction performance. Another future direction is more of the metabolite set tests (or pathways analysis) according to the predicted metabolites.

As a summary, we anticipate the newly developed ENVIM method for microbiome-based metabolite prediction provides good prediction performance and will be used to infer individual metabolites experimental design when only microbiome data are available, or in the condition that a proportion of samples in a study have no metabolome profile.

## Acknowledgments

We acknowledge NIH/NIDCR R03-DE02898, NIH/NIDCR U01-DE025046 and P30 CA016059 (Massey Cancer Center Support Grant) for funding support. We also acknowledge that Prof William Valdar for supporting Jialiu Xie.

## Contribution to the Field Statement

Microbiome data are becoming increasingly available in large health cohorts yet metabolomics data are still scant. While many studies generate microbiome data, they lack matched metabolomics data or have considerable missing proportions of metabolites. Since metabolomics is key to understanding microbial and general biological activities, the possibility of imputing individual metabolites from microbial taxonomy or metagenomics is intriguing. Importantly, current metabolomics profiling methods have unknown accuracy and are limited in their ability to predict individual metabolites. To address this gap, we developed a novel metabolite prediction method (ENVIM) based on the Elastic Net Model (ENM) using metagenomics, metatranscriptomics, or both data types. ENVIM introduces an extra step to ENM to consider variable importance scores and thus achieve better prediction power. The better prediction capability of ENVIM than the existing MelonnPan in three datasets generated from oral, gut, and vaginal samples, suggest the potential usage in a variaty of studies from different body sites. As a summary, we anticipate ENVIM provides good prediction performance, and will be used to infer individual metabolites experimental design when only microbiome data are available, or in the condition that a proportion of samples in a study have no metabolome data profile.

## Figure and Table legend for supplemental files

**Supplemental Figure 1**. Boxplot of -log10 of mean square error for DNA, RNA, and Both in each of the three studies. None of the nominal p values to compare ENVIM and MelonnPan are significant as all of them >0.1.

**Supplemental Figure 2**. Diagnosis for outlier samples. The X-axis is the cumulative proportion of samples, and the y-axis is number of non-missing values. The left lower tail dots that are far from the rest may be considered as sample outliers. For ZOE 2.0 data and Lloyd-Price data, we need to remove the 10 outliers subjects from ZOE 2.0 data and 15 outliers from Lloyd-Price data to ensure the distribution of non-missing values is continuous.

**Supplemental Table 1**. Overall prediction results, for all gene family data types, all three datasets, and both methods, in Spearman correlation and MSE.

**Supplemental Table 2.** The gene lists in DNA or RNA, based on the highest rank or the average rank among metabolites, that contribute to metabolite prediction in ZOE 2.0 by ENVIM. Rank is based on the weight matrix in ENVIM. A larger number of ranks suggests more important gene families.

