## Supplementary figures and images for "Improved Metabolite Prediction Using Microbiome Data-Based Elastic Net Models"

### Supplemental Figure 1

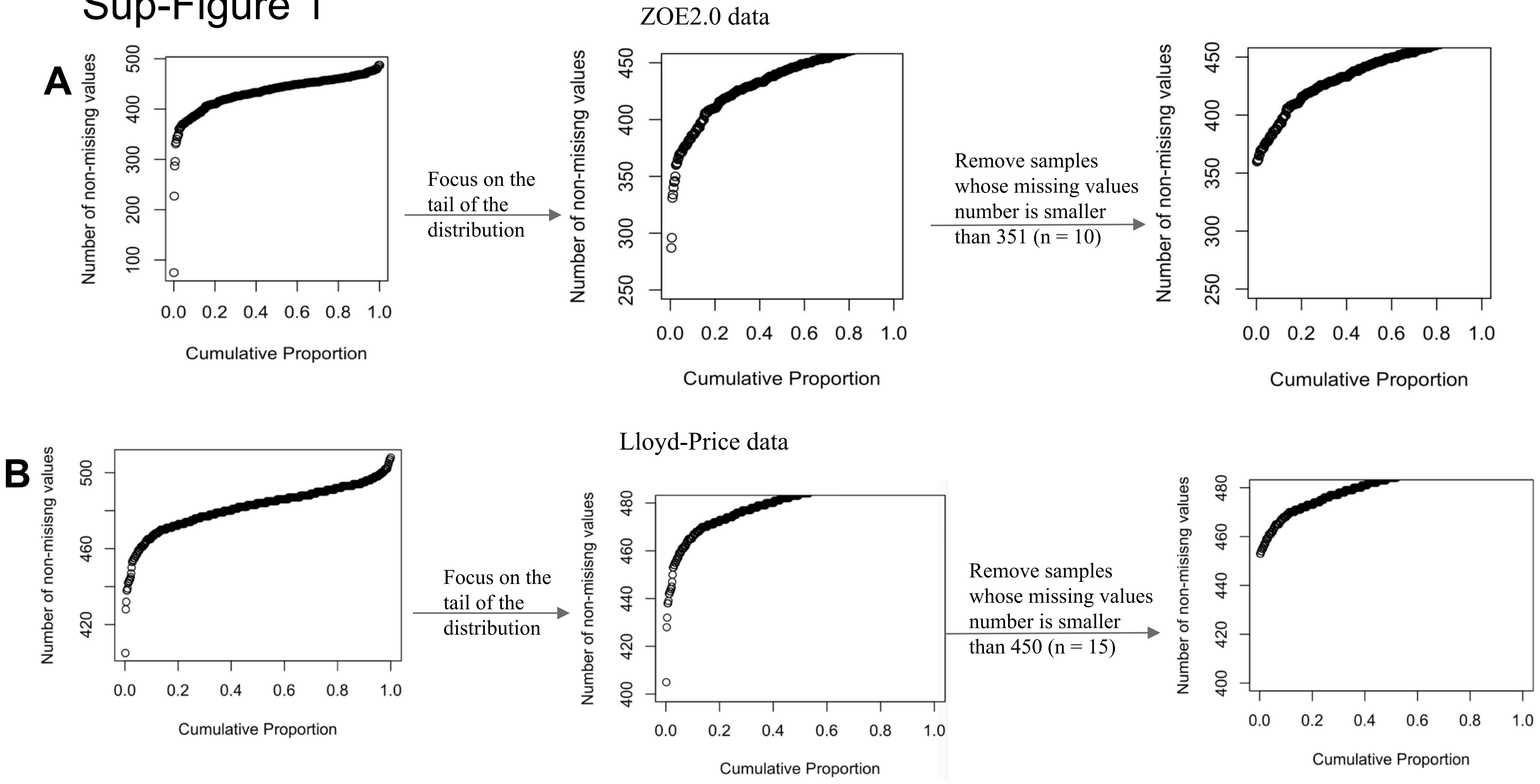

### Supplemental Figure 2

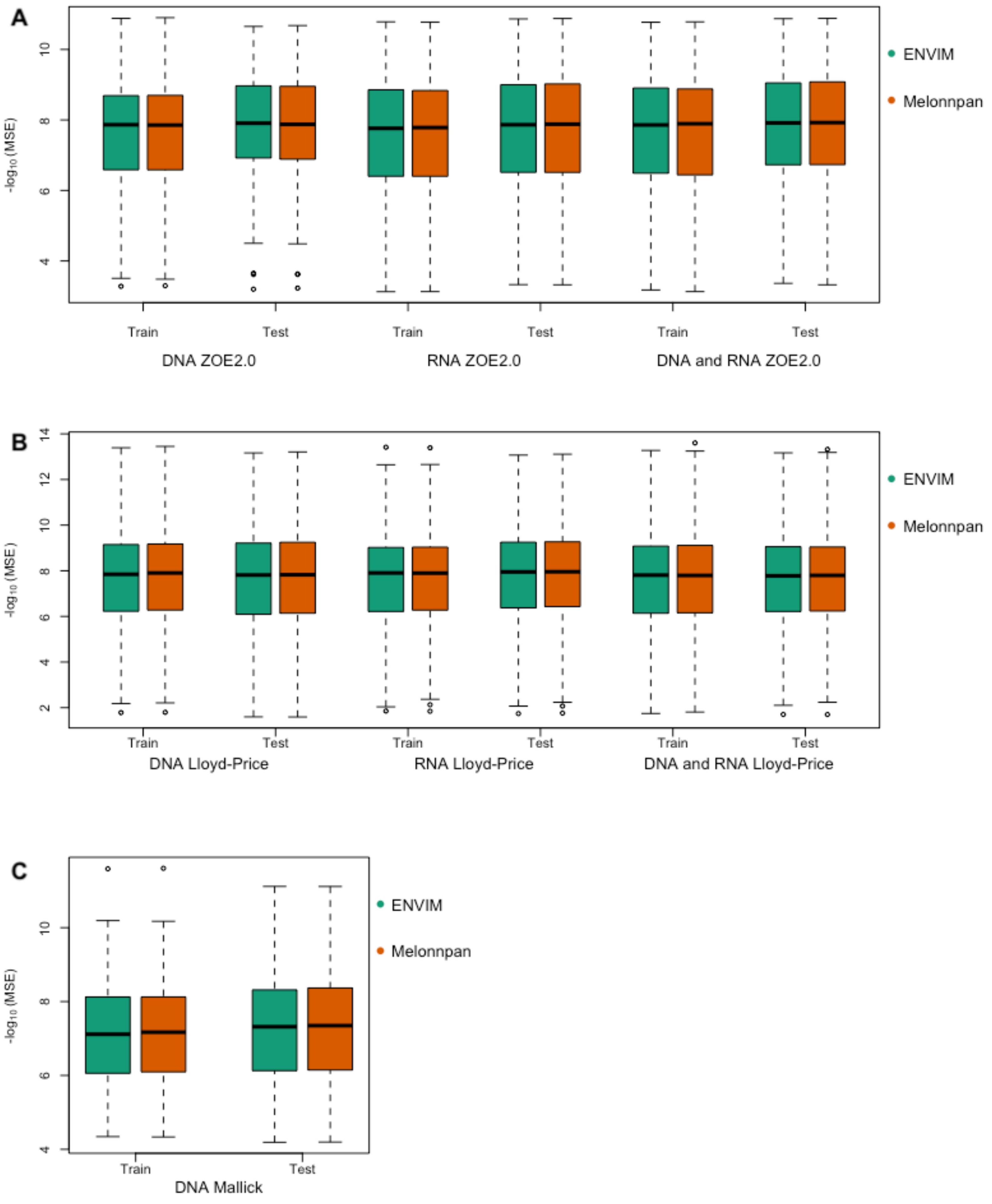
